# Cenote-Taker 3 for Fast and Accurate Virus Discovery and Annotation of the Virome

**DOI:** 10.1101/2025.08.20.671380

**Authors:** Michael J. Tisza, Arvind Varsani, Joseph F. Petrosino, Sara J. Javornik Cregeen

## Abstract

Viruses are abundant across all Earth’s environments and infect all classes of cellular life. Despite this, viruses are something of a black box for genomics scientists. Their genetic diversity is greater than all other lifeforms combined, their genomes are often overlooked in sequencing datasets, they encode polyproteins, and no function can be inferred for a large majority of their encoded proteins. For these reasons, scientists need robust, performant, well-documented, extensible tools that can be deployed to conduct sensitive and specific analyses of sequencing data to discover virus genomes - even those with high divergence from known references - and annotate their genes. Here, we present Cenote-Taker 3. This command line interface tool processes genome assemblies and/or metagenomic assemblies with modules for virus discovery, prophage extraction, and annotation of genes and other genetic features. Benchmarks show that Cenote-Taker 3 outperforms most tools for virus gene annotation in both speed (wall time) and accuracy. For virus discovery benchmarks, Cenote-Taker 3 performs well compared to geNomad, and these tools produce complementary results. Cenote-Taker 3 is freely available on Bioconda, and its open-source code is maintained on GitHub (https://github.com/mtisza1/Cenote-Taker3).

## Introduction

Cataloging the virome from *de novo* assembled sequencing data remains a primary challenge in the field of virus metagenomics. Essential subtasks within this challenge include identifying virus genomes over viral-like genetic elements in host genomes, taxonomically classifying them, functionally annotating their genes, and extracting prophages (integrated virus genomes in prokaryotic host genomes) and proviruses (viruses in eukaryotic chromosomes) via virus/cellular genome boundary^1^. Until bioinformatics tools can sufficiently conduct these tasks, downstream analyses such as association of viruses with specific host phenotypes (e.g. health vs. disease), deep understanding of virus evolution through comparative genomics, and microbial ecology of viruses and their hosts all remain underwhelming and subject to errors and biases in catalog composition and annotation.

Virus discovery from metagenomic assemblies is commonly performed with tools such as VirSorter2^2^, VIBRANT^3^, and geNomad^4^, which use combinations of hallmark-gene detection and (in some cases) machine-learning classifiers to distinguish viral from cellular or plasmid sequences. For functional annotation and genome interpretation, many workflows rely on ORF calling with prodigal^5^, prodigal-gv (https://github.com/apcamargo/prodigal-gv), or PHANOTATE^6^ and phage-focused functional annotators such as Pharokka^7^ (PHROGs-based), broader annotation frameworks such as MetaCerberus^8^, and complementary approaches such as phold^9^ that leverage structure-informed protein annotation. Downstream quality control utilities like CheckV^1^ are used for completeness/contamination estimates, and specialty tools like CRESSENT^10^ can perform value-added analyses on certain taxa (e.g. CRESS-DNA viruses). Despite having various software packages for a complete viromics “workflow”, major gaps remain in the discovery and characterization genetic entities that are extremely divergent from well-characterized records.

Cenote-Taker 3 takes on these challenges to advance viral genomics for discovery and annotation of virus genomes from *de novo* assembled sequences (contigs). This tool scales from a single genome to terabase-size datasets. The complete Cenote-Taker 3 workflow can be summarized as follows: 1) predict and translate open reading frames from input contigs, 2) detect presence/absence of viral hallmark genes in each contig to detect putative virus contigs, 3) explore these contigs for terminal repeats or circularity, wrapping and rotating potentially circular contigs, 4) annotate genes by function, 5) extract prophages in contigs with high bacterial gene content, and 6) assign hierarchical taxonomy labels to each virus sequence (Fig. 1). Output files include gene- and contig-level summary tables, FASTA format files for nucleotide and protein sequences, and GenBank flat files as interactive genome maps for each predicted virus. In this manuscript, we benchmark Cenote-Taker 3 against several of the most prominent virus metagenomics tools to determine its utility in gene annotation, virus discovery, and scalability.

**Figure 1.**
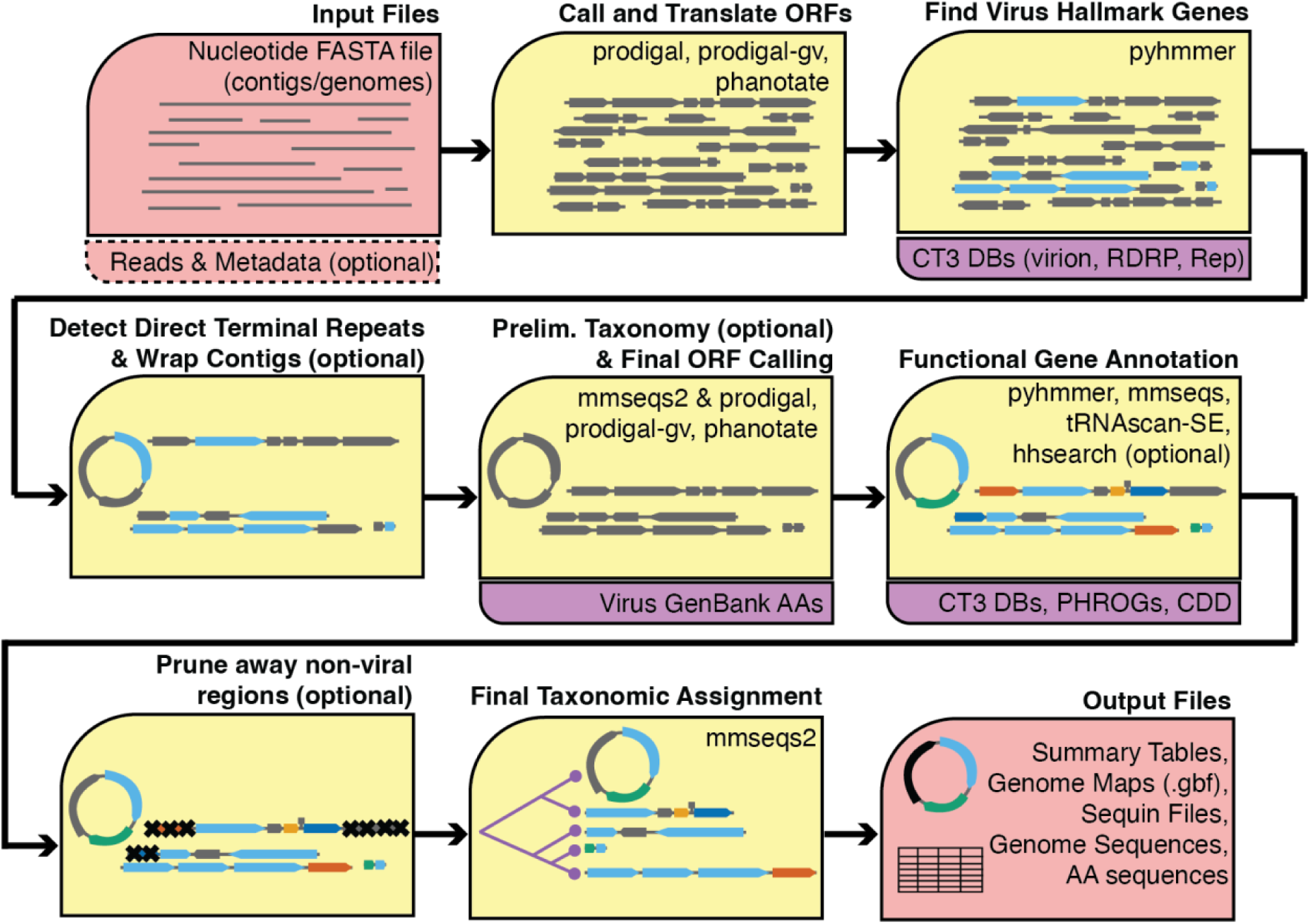
Cenote-Taker 3 Schematic. A visual description of how input contigs are systematically processed by Cenote-Taker 3 to identify virus genomes, produce gene annotations, prune prophages, and assign taxonomy. Output files include genome maps (.gbf), tabular virus- and gene-level summaries, and sequences (.fasta).

Benchmarking of bioinformatics software, while essential, can be difficult to interpret due to choices made and the priorities set by the individuals conducting the benchmarking tests. One approach in virus prediction benchmarks is to synthesize ever smaller snippets of nucleotide sequences of known viruses and then determine if these snippets can be recalled by virus prediction software^11^. While this may have some use, we argue that, for the purposes of cataloging the virome, these tests are of minimal utility, as they do not measure the ability of software to identify previously undiscovered, highly divergent virus genomes. Furthermore, contigs representing complete or mostly complete virus genomes with identifiable coding regions / functional domains are substantially more valuable and informative to catalogs than incomplete genome fragments.

In the future, we envision biome-specific catalogs of complete, well-characterized virus genomes that can be used in quantitative read-mapping analyses, comparative genomics and other biotechnology applications. To achieve this, we consider what different technologies can offer. Metagenomic assembly algorithms utilizing short reads (produced by the standard technologies for over a decade) produce a plurality of short contigs usually representing small genome fragments. However, long read technologies, such as those employed by PacBio and Oxford Nanopore Technologies (and under development by other companies), produce reads from metagenomic libraries of sufficient quantity, quality, and length to allow assembly of complete cellular (e.g. bacterial), plasmid, and virus genomes. And these technologies are increasingly applied to metagenomes^12^.

Therefore, because of their utility and increasing obtainability, the benchmarks herein focus on single-contig MAGs (metagenome-assembled genomes) representing complete or high-quality genomes from a diversity of ecosystems. The virus genomes in these benchmarks encoding genes that primarily have no or little amino acid similarity to protein records in the large public RefSeq Virus repository^13^.

All things considered, Cenote-Taker 3 is a high-performing tool that allows researchers to discover and annotate the virome in complex sequence datasets.

## Results

### Improvements Over Cenote-Taker 2

Cenote-Taker 3, while sharing the same goals as its predecessor^14^, completely eclipses its utility and performance. The codebase has been completely rewritten, additional utilities were added, it is much more efficient (5-fold decrease in wall time for the same dataset; Fig. S1B), the database is greatly expanded and more consistently annotated (Fig. S1A). Also, it is now available via the Bioconda package manager^11^.

**Figure S1.**
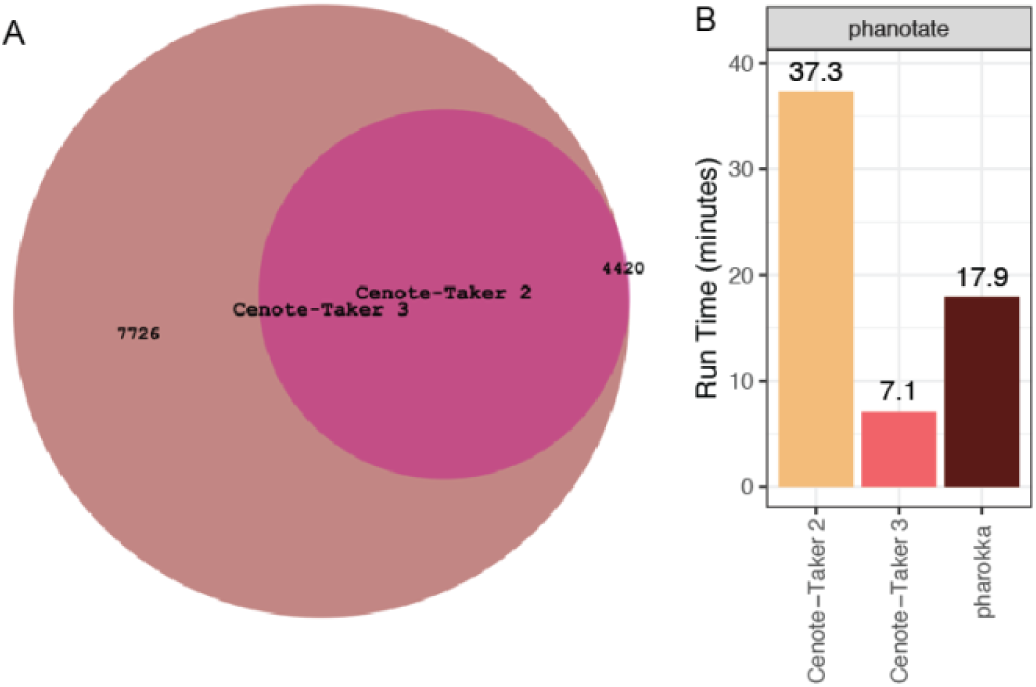
Updates from Cenote-Taker 2. (A) The hallmark gene Hidden Markov Model database from Cenote-Taker 2 was expanded for Cenote-Taker 3 by adding 7,726 new models. (B) Comparing time to process 100 UHGV genomes between Cenote-Taker 2, Cenote-Taker 3, and Pharokka. The phanotate open reading frame predictor was used as it is the only setting that is comparable to Cenote-Taker 3.

### Functional Gene Annotation

Cenote-Taker 3 exists in a rich ecosystem of bioinformatics tools that are purported to or can be used to annotate genes and other features on virus genomes. Not all software packages are tested here. While Cenote-Taker 3 has the capability (because of the scope of its curated databases) to annotate (and discover) viruses-infecting all domains of life, most other tools focus on annotation of bacteriophage genomes, which, via their ubiquity and immense diversity, provide ample test sets.

The UHGV catalog^15^ has compiled hundreds of thousands of virus MAGs of varying levels of completeness, and UHGV primarily consists of dsDNA and ssDNA phages from human guts. Here we have taken random subsets of 100 and 1,000 high-quality virus MAGs (Table 1) and compared to state-of-the art tools geNomad^4^ (mmseqs-based), MetaCerberus^8^ (hmmer-based), Pharokka^7^ (hmmer-based), and phold^9^ (foldseek-based^16^), and Cenote-Taker 3 (hmmer- and mmseqs-based). Here, Cenote-Taker 3 annotates a significantly higher proportion of genes than all tools except phold (Fig. 2A). It has a shorter wall time than all tools except geNomad (Fig. 2B). Note that phold is the only software requiring GPU capabilities for gene annotation from this set. Wall time rankings remain consistent when the dataset is scaled to 1,000 virus MAGs (Fig. S2E).

**Figure 2.**
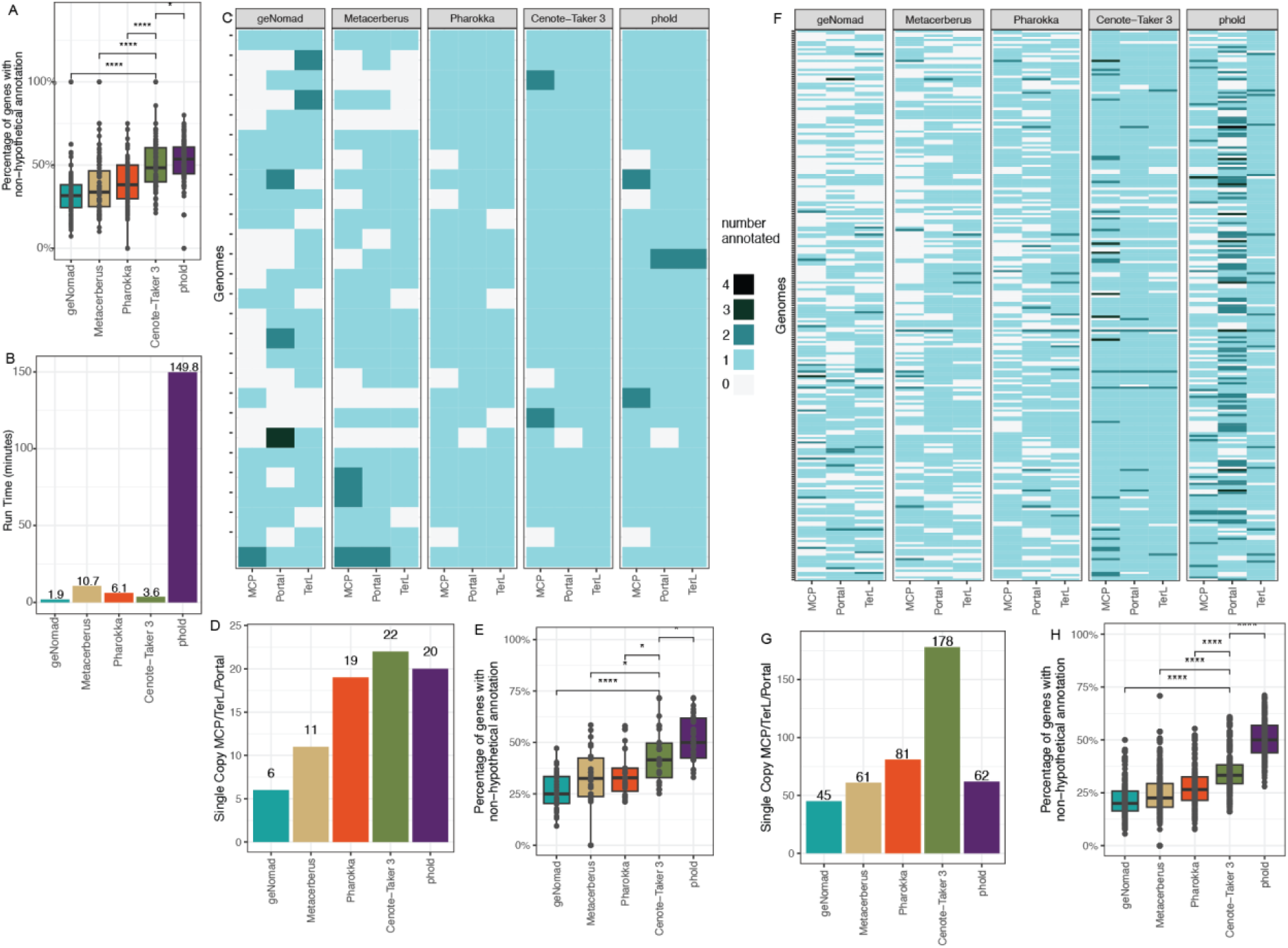
Virus gene annotation benchmarks. (A) UHGV 100 virus Metagenome-Assembled Genomes (MAGs) annotated with five different software packages. The y-axis reports annotated (non-hypothetical) genes as a percentage of total genes. (B) Wall time for UHGV 100 virus MAGs annotation. (C) 27 circular gut (stool)-derived head-tail phage MAGs annotated by five software packages. Per-genome number of annotated major capsid protein (MCP), large terminase subunit (TerL), and portal protein genes. (D) Out of 27 circular gut (stool)-derived head-tail phage MAGs, number of genomes with single copy MCP/TerL/portal genes. (E) 27 circular gut (stool) head-tail phage MAGs, reporting annotated (non-hypothetical) genes as a percentage of total genes. (F) Like (C) but with 242 seawater-derived head-tail phage MAGs. (G) Like (D) but with 242 seawater-derived head-tail phage MAGs. (H) Like (E) but with 242 seawater-derived head-tail phage MAGs.

**Figure S2.**
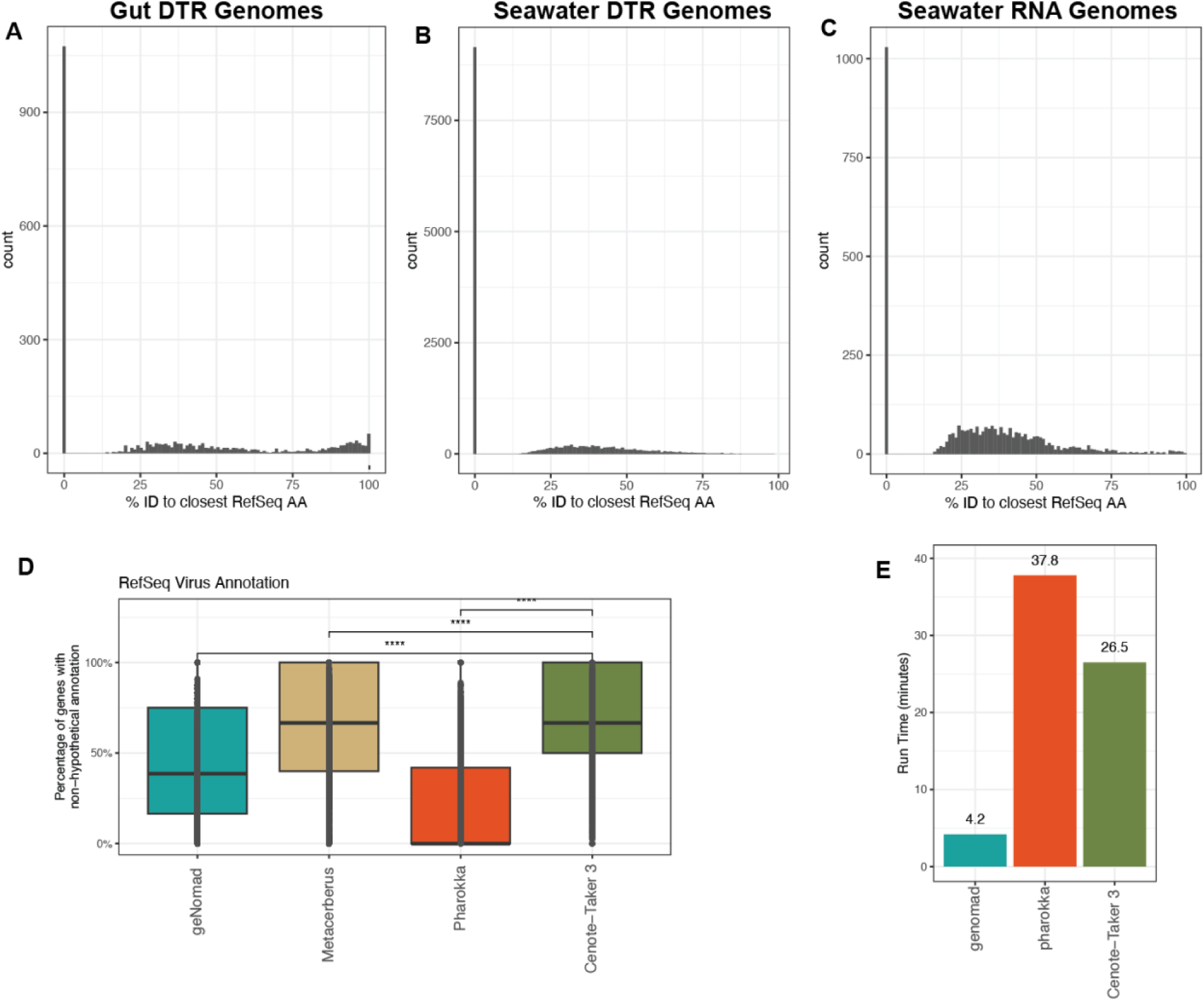
Additional Annotation Analyses. Translated amino acid sequences were compared to RefSeq Virus amino acid sequences using mmseqs (e-value cutoff 0.01) for (A) Gut DTR Genomes, (B) Seawater DTR Genomes, and (C) Seawater RNA Genomes. (D) RefSeq Virus genomes (n=18,969 contigs) annotated with four different software packages. The y-axis reports annotated (non-hypothetical) genes as a percentage of total genes. (E) Wall Time for Annotating 1,000 UHGV Genomes.

**Table 1:**
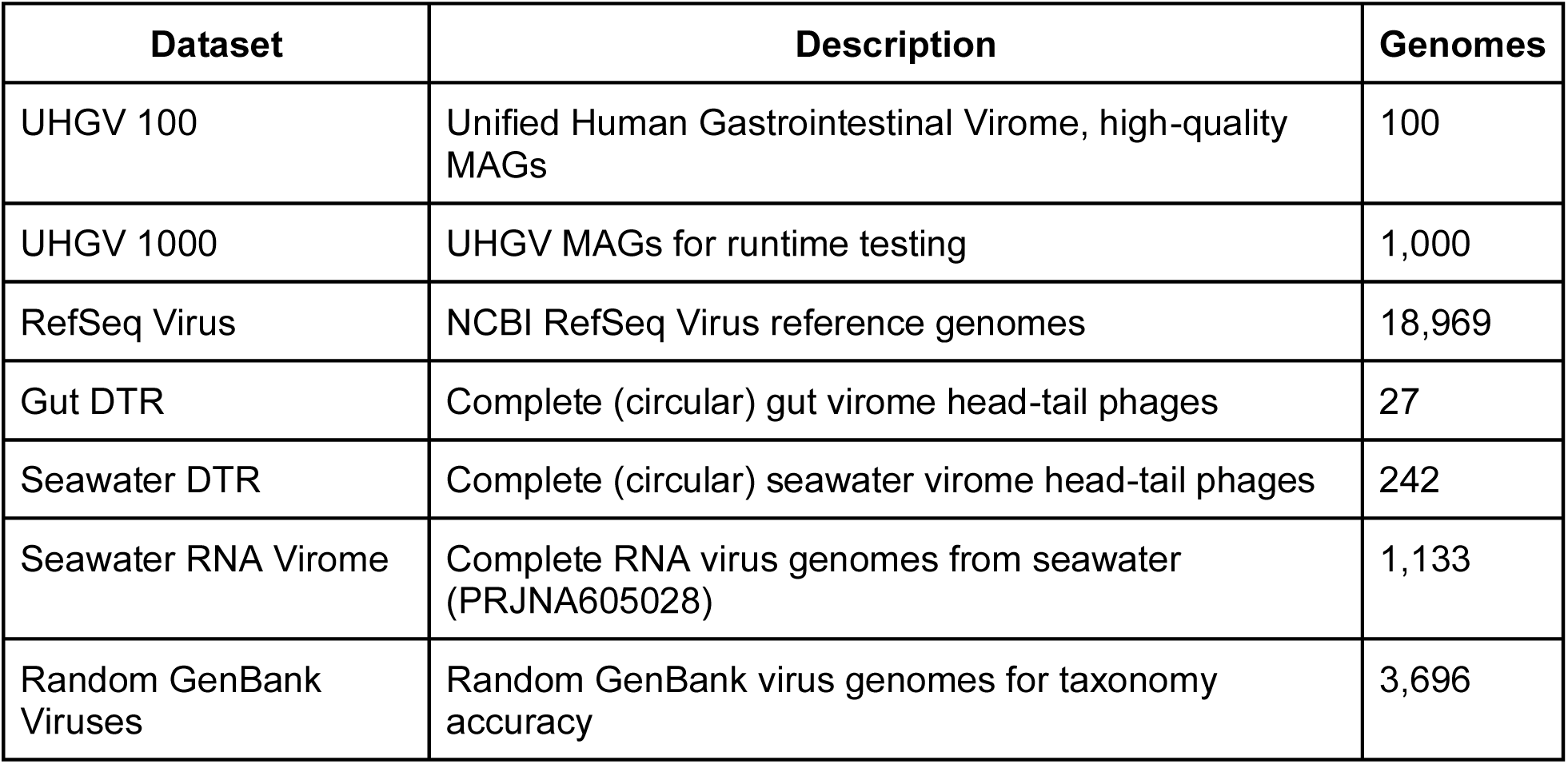
Annotation Datasets.

Simply measuring the proportion of genes annotated does not account for the possibility of false positives or improper labeling. Considering the test data are mostly divergent genomes (Fig. S2A-C) that are difficult to assess for ground truth, we reasoned that some features should be nearly universal in some virus categories. For example, head-tail phages (viruses in the class *Caudoviricetes*) should have one and only one copy of a major capsid protein (MCP) gene, a large terminase subunit (TerL) gene, and a portal protein gene. Software packages can be benchmarked on their ability to annotate a single copy for each of these conserved genes. Therefore, suitable datasets of putatively complete *Caudoviricetes* genomes were needed to assess specific marker gene annotation.

A recent study used long read data plus short read data to assemble and polish contigs from the viromes of human gut (stool) and seawater virus-like particle preps ^17^ (Table 1). These assemblies contained many complete head-tail phages, and we used these genomes to assess annotation accuracy of major capsid (MCP), large terminase (TerL), and portal genes. In both the gut dataset (Fig 2C-E) and the seawater dataset (Fig2F-H), Cenote-Taker 3 annotates the most head-tail phage genomes with correct number of these three genes. Figure 3 C and F demonstrate that all tools are susceptible to missing these genes (false negatives) as well as predicting multiple copies of the target genes (false positives). Cenote-Taker 3 annotated the expected number of MCP/TerL/Portal protein genes in 22/27 head-tail phages (81.4% perfect rate) in the gut dataset surpassing geNomad (22.2%), MetaCerberus (40.7%), Pharokka (70.3%), and phold (74.1%) (Fig. 2D). And Cenote-Taker 3 correctly classified 174/242 (73.5% perfect rate) in the seawater dataset, outpacing geNomad (18.5%), MetaCerberus (25.2%), Pharokka (33.5%), and phold (25.6%) (Fig. 2G). Additional statistics are available in Supplementary Tables S1-6. The combination of speed, annotation rate, and accuracy demonstrate that Cenote-Taker 3 is a strong choice for gene annotation tasks for virus MAGs.

**Figure 3.**
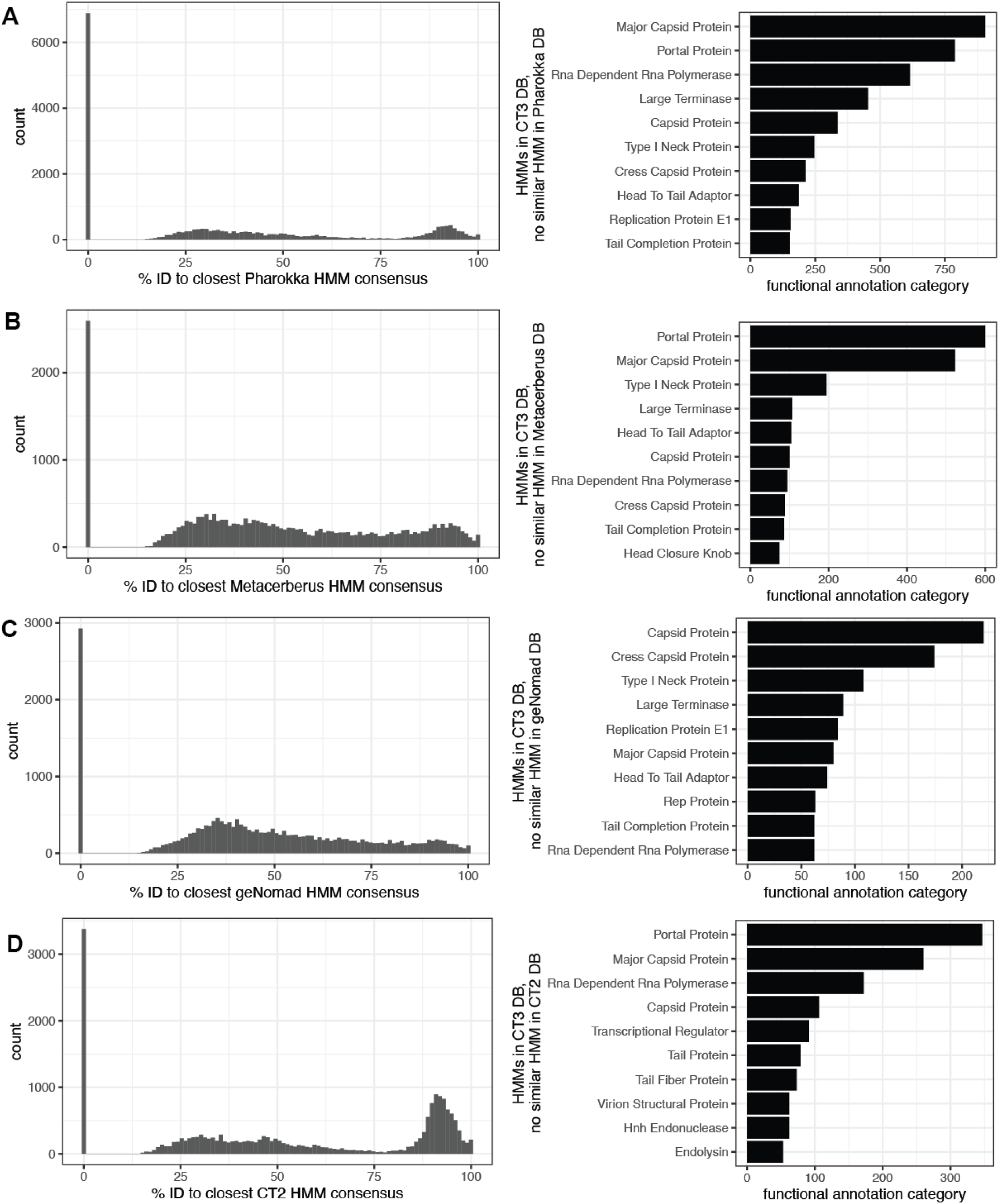
Comparison of Gene Family HMMs. Consensus sequences from all individual HMMs were extracted from Cenote-Taker 3, Pharokka, MetaCerberus, geNomad and Cenote-Taker 2 and compared with mmseqs2. (A) Left: histogram of amino acid identity of consensus sequences of Cenote-Taker 3 DB to Pharokka DB. Right: top 10 functional categories of Cenote-Taker 3 models with no detected similar to Pharokka models. (B) Like A but with MetaCerberus and Cenote-Taker 3. (C) Like A but with geNomad and Cenote-Takaer 3. (D) Like A but with Cenote-Taker 2 and Cenote-Taker 3.

RNA viruses, such as those that might be sequenced in metatranscriptomics studies, have more diverse genome end types and may consist of multiple segments. Therefore, determining completeness of previously undiscovered viruses is more difficult. Nevertheless, other, potentially more biased, methods exist to detect complete virus MAGs. Here, MAGs from an RNA sequencing dataset from a large seawater virus-like particle preparation (Table 1) were filtered using CheckV to retain putatively complete and high-quality virus genomes (Fig. S2C). These genomes were annotated by the same five software packages, and Cenote-Taker 3 had the highest proportion of annotated genes and had the highest number of contigs with an annotated RNA-dependent RNA polymerase (RdRP) gene and Capsid/Coat gene (Fig. S3). It should be noted that Pharokka and phold are designed for DNA phages, and their decreased performance should be understood as a design choice.

**Figure S3.**
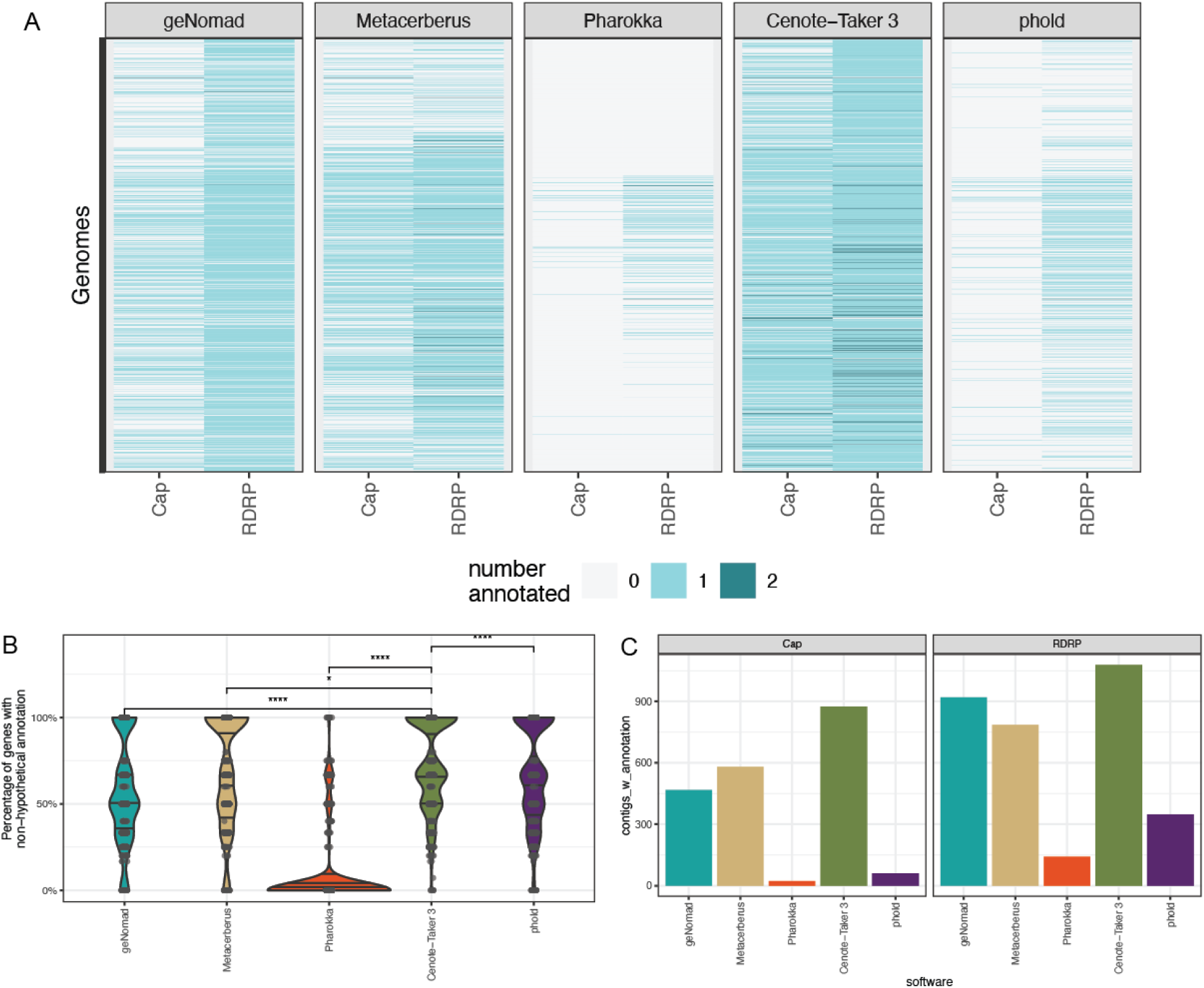
Annotation of RNA virus genomes. (A) 1,133 putatively complete RNA virus MAGs annotated by five software packages. Per-genome number of annotated capsid/coat protein (Cap) and RNA-Dependent RNA Polymerase (RdRP). (B) 1,133 putatively complete RNA virus MAGs annotation rate. The y-axis reports annotated (non-hypothetical) genes as a percentage of total genes. (C) Out of 1,133 putatively complete RNA virus MAGs, number of genomes with Cap and RdRP annotated, respectively.

To compare annotation performance on a well-characterized and diverse dataset, each software package was run on complete RefSeq Virus genomes (n=18,969) (Table 1). In this test, Cenote-Taker 3 annotated the highest proportion of genes on average, followed closely by MetaCerberus (Fig. S2D).

Next, we wanted to resolve why Cenote-Taker 3 outperformed other software packages in these virus annotation benchmarks despite using similar methods (hmmer and mmseqs) as most other approaches. Cenote-Taker 3, Pharokka, MetaCerberus, and Cenote-Taker 2 have associated HMM databases containing models for virus gene families, and these databases could be compared. Consensus amino acid sequences of all models were extracted and aligned using mmseqs2^18^ (e-value threshold 0.01). Across all comparisons, Cenote-Taker 3’s database contained thousands of models with no similarity to models in other databases (Fig. 3A-D). Importantly, when we looked at the top 10 functional labels in Cenote-Taker 3-specific models (i.e. no mmseqs alignments to models in other tools), we see that models for Major Capsid Protein, Portal protein, Large Terminase, RNA-Dependent RNA Polymerase, and other key hallmark genes are the most abundant. Therefore, it’s almost certain that the gene annotation improvements are primarily a function of more and better gene family models utilized by Cenote-Taker 3.

### Virus Discovery

To assess virus discovery capabilities, we compared Cenote-Taker 3 with geNomad on circular contigs from two distinct environmental datasets: hot springs^19^ and anaerobic digester metagenomes^20^ (Table 2). GeNomad was chosen since it has become a dominant virus discovery software in recent years and is well-maintained. It uses a combination of marker gene annotation and *k*mer-based neural network classification to predict which contigs are viruses. Cenote-Taker 3 uses a solely marker gene-based approach to virus discovery.

**Table 2:** Discovery Datasets.

| Dataset | Description | Size | Sequences |
| --- | --- | --- | --- |
| DRR290133 | Long-read assembly, circular contigs | 0.168 Gb | 301 |
| ERR10905741 | Long-read assembly, circular contigs | 0.175 Gb | 390 |
| ERR8081307 | Short-read VLP assembly | 0.014 Gb | 5,666 |
| ERR8081362 | Short-read VLP assembly | 0.007 Gb | 2,699 |
| ERR8081370 | Short-read VLP assembly | 0.006 Gb | 3,673 |
| ERR13455686 | Short-read bulk WGS assembly | 0.018 GB | 4,909 |
| SRR36807671 | Short-read bulk WGS assembly | 0.032 Gb | 16,397 |
| SRR36931851 | Short-read bulk WGS assembly | 0.18 Gb | 67,543 |

Long read datasets were assembled with myloasm v0.1.0^21^ and filtered for circularity. Circular contigs should represent either complete bacterial chromosomes (which may contain prophages), plasmids, or circular DNA virus genomes. These were run through Cenote-Taker 3 and geNomad using virus discovery/detection settings recommended in each software’s documentation.

Overall, the two methods largely agree on the hot springs data (Fig. 4A). However, there is some disagreement. Again, ground truth is difficult if not impossible to know when working with this data type. To analyze Cenote-Taker 3-specific and geNomad-specific contigs, phold was used to annotate genes since this method’s structure-based approach is somewhat orthogonal to other methods. We find that both discovery methods missed contigs with complete arrays of major capsid protein, large terminase subunit, and portal genes (Fig. 4B-C). Inspection of phold-based genome maps is interesting if not conclusive (Fig. 4D,E). Many Cenote-Taker 3-specific contigs encode high numbers of phage tail-related genes, for example, which may represent phage-derived tailocin elements.

**Figure 4.**
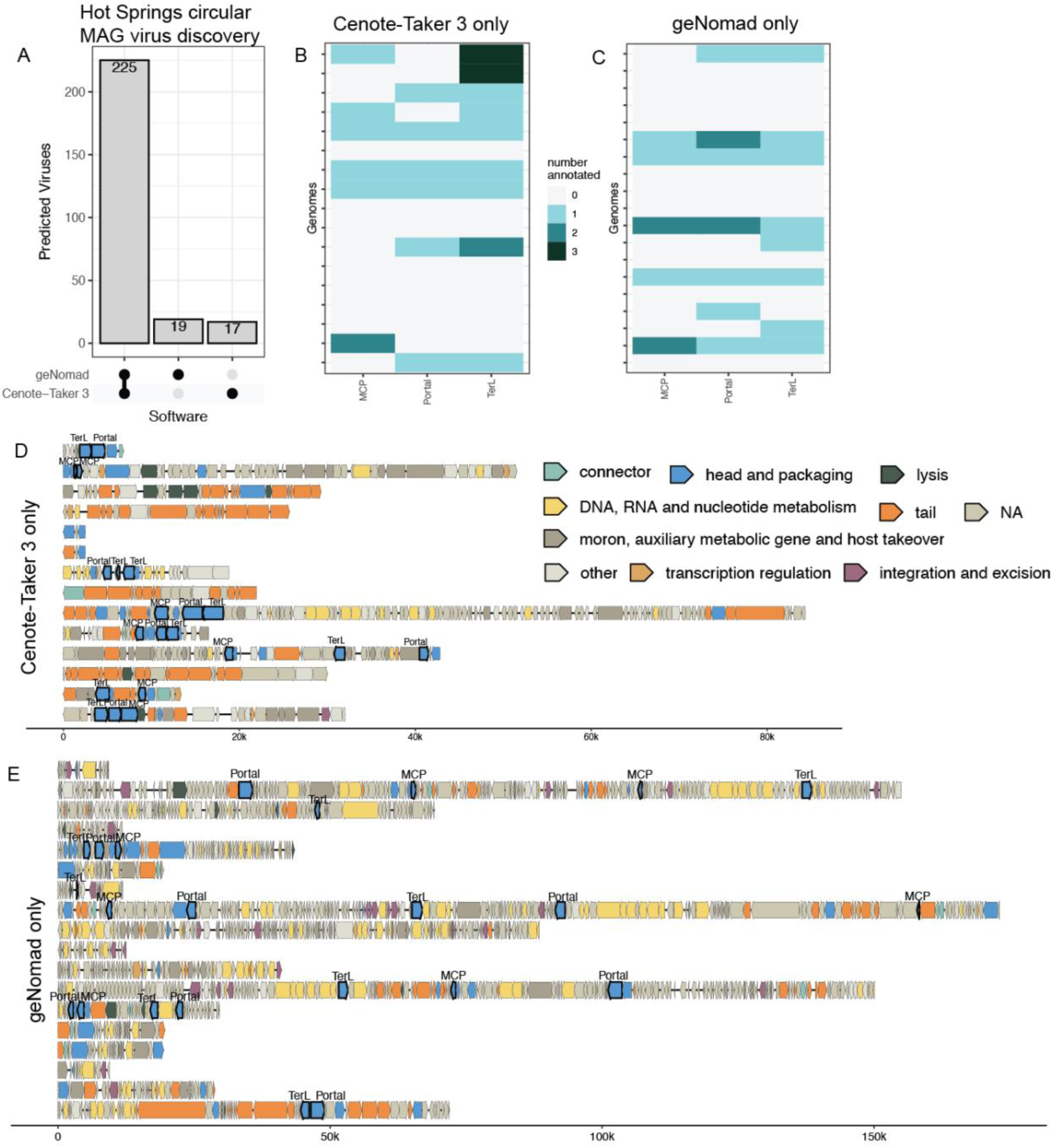
Virus Discovery Comparison for Hot Springs Data. (A) Comparison of viruses identified/discovered between geNomad and Cenote-Taker 3 in a dataset of circular MAGs from a hot spring long read metagenome assembly. (B) For viruses in this dataset only identified by Cenote-Taker 3, count of MCP/TerL/Portal protein genes annotated per contig by phold. (C) Like (B) but for viruses in this dataset only identified by geNomad. (D) For viruses in this dataset only identified by Cenote-Taker 3, genome maps of phold annotations, highlighting MCP/TerL/Portal protein genes. Only contigs with under 200 genes shown for readability. (E) Like (D) but for viruses in this dataset only identified by geNomad.

Results were more divergent in the anaerobic digester dataset (Fig. 5A), wherein geNomad called 143 unique contigs while Cenote-Taker 3 called 37 unique contigs viral. We find that Cenote-Taker 3 has a greater number of unique contigs that encode major capsid protein, large terminase, and/or portal protein genes, per phold annotation (Fig. 5B,D). GeNomad, on the other hand, has many unique contigs from the anaerobic digester dataset that have few of these marker genes (despite geNomad taxonomic labels of *Caudoviricetes* for nearly all contigs), and, interestingly, most are about 25 kilobases in length (Fig. 5C,E). The geNomad output files show that the large majority of the unique contigs encoded no hallmark genes (109/143) and the software used predictions from its neural network on these MAGs (Fig. S4). Shared gene content (not shown) suggests these contigs may represent a coherent class of elements. It’s hard to say whether these elements are truly viruses, but it would also be difficult to ascribe any other category to them.

**Figure 5.**
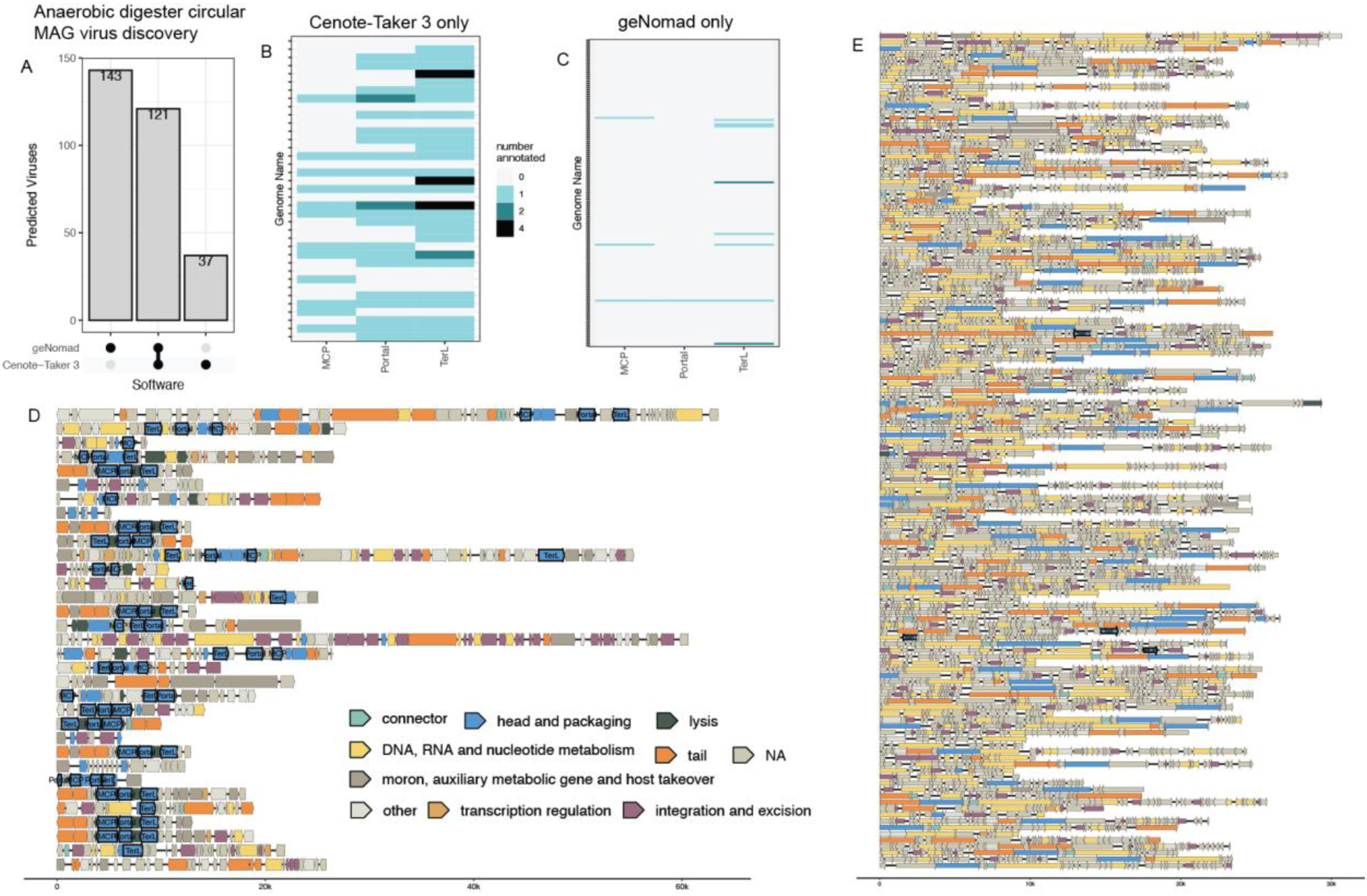
Virus Discovery Comparison for Anaerobic Digester Data. (A) Comparison of viruses identified/discovered between geNomad and Cenote-Taker 3 in a dataset of circular MAGs from anaerobic digester long read metagenome assembly. (B) For viruses in this dataset only identified by Cenote-Taker 3, count of MCP/TerL/Portal protein genes annotated per contig by phold. (C) Like (B) but for viruses in this dataset only identified by geNomad. (D) For viruses in this dataset only identified by Cenote-Taker 3, genome maps of phold annotations, highlighting MCP/TerL/Portal protein genes. Only contigs with under 200 genes shown for readability. (E) Like (D) but for viruses in this dataset only identified by geNomad.

While we do not claim that Cenote-Taker 3 is the best approach to use when searching metagenomic data for highly fragmented and incomplete virus genomes, it is nevertheless informative to see how it performs in virus discovery tasks on short-read metagenomics assemblies. Six datasets from gut (stool) DNA – three from bulk WGS and three from virus-like particle preps – were assembled (Table 2). Then, Cenote-Taker 3 and geNomad were run in discovery mode, using a two marker/hallmark gene minimum to avoid false positives. While no ground truth data is available to validate these benchmarks, the union between these tools is greater than the differences (Fig. S5), suggesting that either software might be used for this task. Further, using different parameters and thresholds will affect the results.

When these results are taken together, potential users of these software packages face a difficult choice. Using Cenote-Taker 3, putative virus MAGs may be easier to verify since presence of virus hallmark genes can be checked with orthogonal methods. With geNomad, the virus MAGs primarily predicted by the neural network are harder to verify but may contain genomes from entirely undescribed types of viruses.

**Figure S4.**
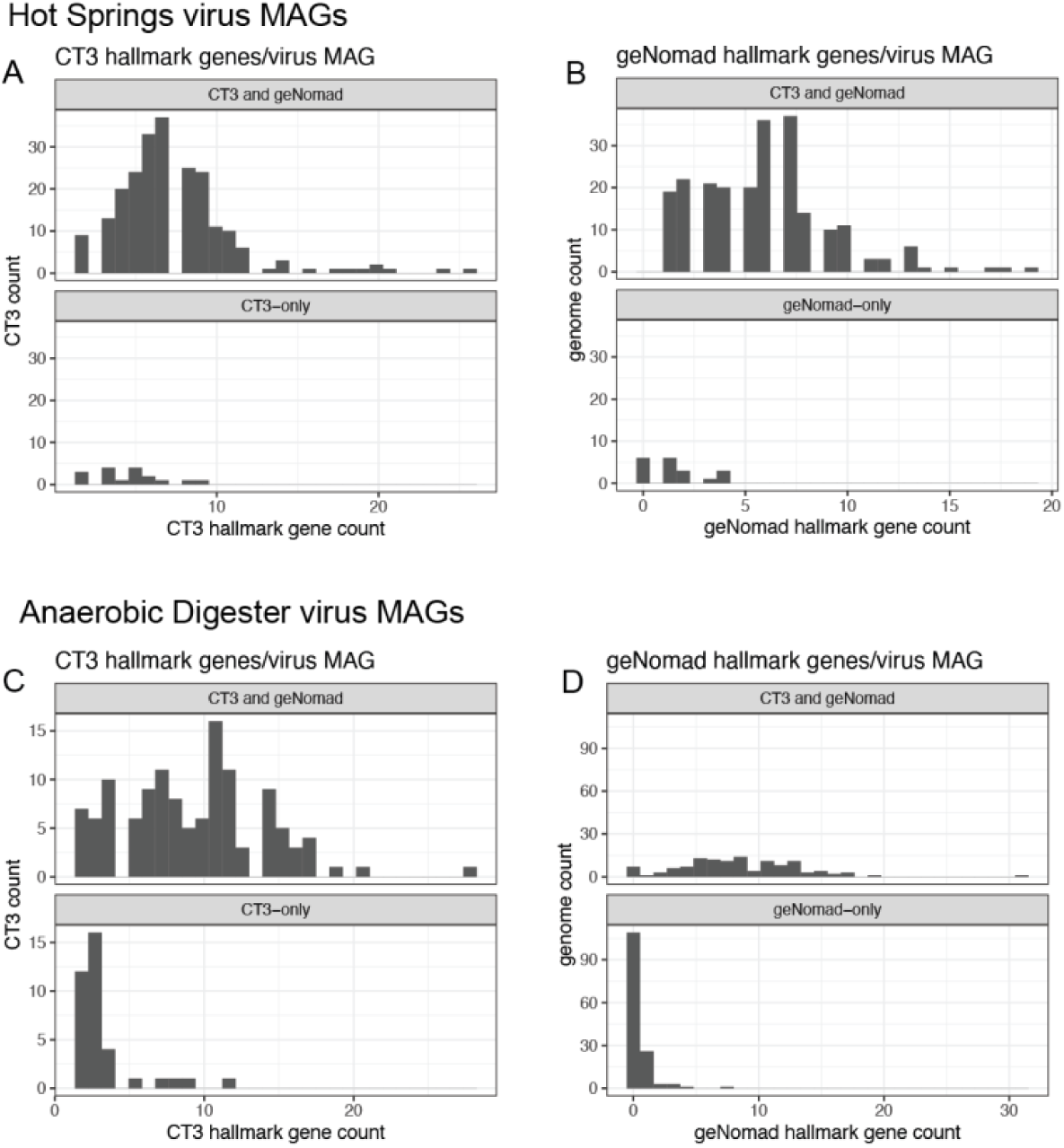
Virus MAG Hallmark Comparison for Shared and Unique Detections. (A) Comparison of hallmark genes for viruses identified/discovered between geNomad and Cenote-Taker 3 in a dataset of circular MAGs from hot spring long read metagenome assembly. Cenote-Taker 3 hallmark gene counts. (B) Like (A), but geNomad hallmark gene counts. (C) Like (A) but for circular MAGs from anaerobic digester long read metagenome assembly. Cenote-Taker 3 hallmark gene counts. (D) Like (C) but geNomad hallmark gene counts.

**Figure S5.**
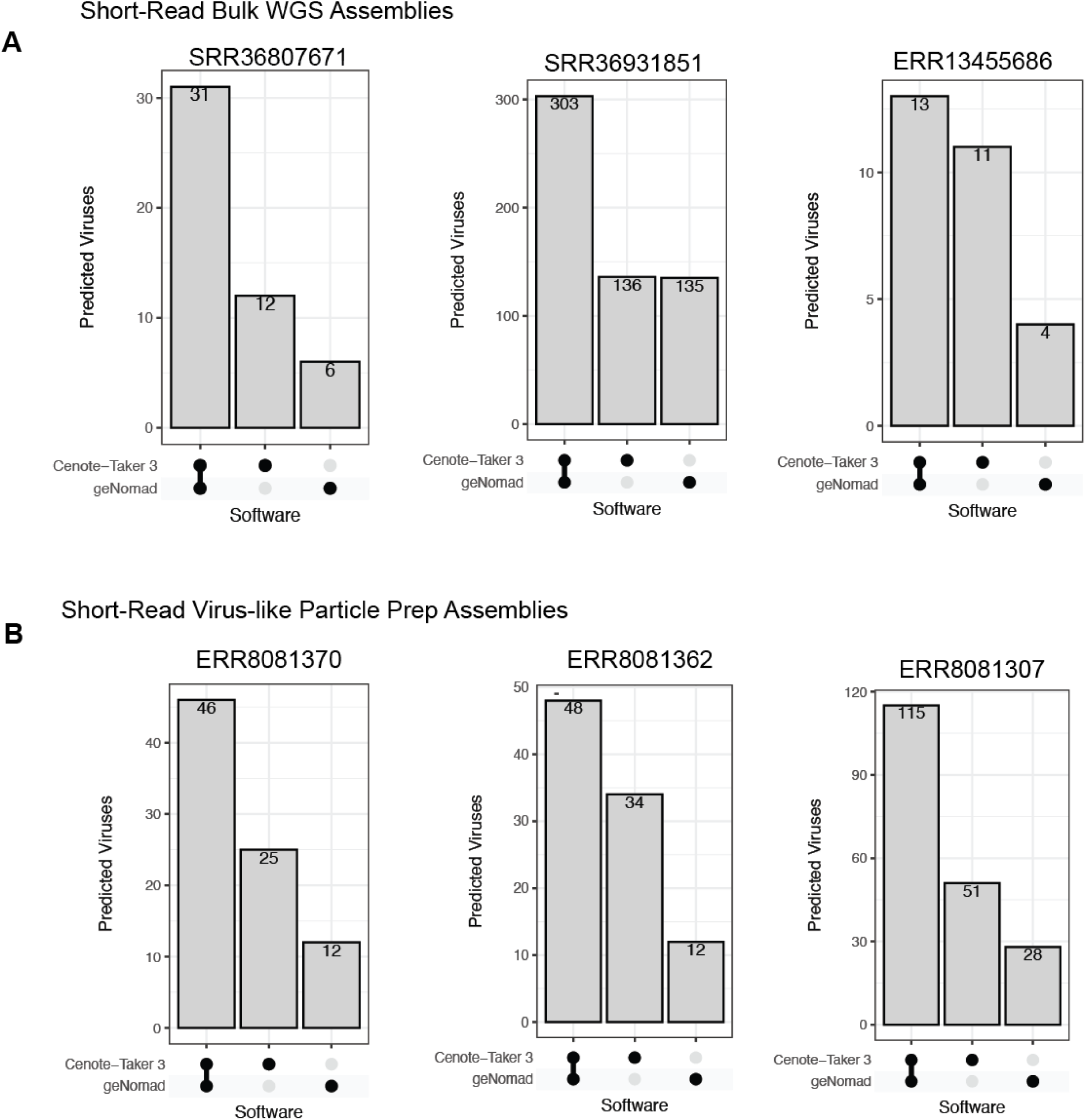
Virus Discovery on Short-Read Assemblies. (A) Three bulk WGS sequencing runs for stool samples were downloaded from SRA and assembled. Cenote-Taker 3 and geNomad were run with comparable settings, and specific contigs that are predicted to be viruses are compared. (B) Like A but with Virus-like Particle preps from stool samples.

### Computational Performance and Resource Scaling

In an era where large metagenomic studies often produce tens to hundreds of gigabases (Gb) of metagenomic contigs, it is important to understand how your software of choice scales. Sampling a large pool of metagenomic contigs at 3% (0.15 Gb), 10% (0.49 Gb), 30% (1.50 Gb), and 100% (5.18 Gb) (Table 3), we compared the wall time and random-access memory usage of Cenote-Taker 3 and geNomad on compute nodes with 1 to 32 CPUs (central processing units).

**Table 3:** Resource Scaling Datasets.

| Dataset | Subset | Size | Sequences |
| --- | --- | --- | --- |
| Large Metagenome (DRR582205 + ERR9769281 + ERR10905741) | 100% | 5.18 Gb | 177,667 |
| Large Metagenome (DRR582205 + ERR9769281 + ERR10905741) | 30% | 1.50 Gb | 53,667 |
| Large Metagenome (DRR582205 + ERR9769281 + ERR10905741) | 10% | 0.49 Gb | 17,876 |
| Large Metagenome (DRR582205 + ERR9769281 + ERR10905741) | 3% | 0.15 Gb | 5,417 |

This benchmark demonstrates that both Cenote-Taker 3 and geNomad can handle very large individual datasets with these resources. Cenote-Taker 3 was faster (higher throughput) when using 1 or 2 CPUs, there were data-dependent results for 4 or 8 CPUs, and geNomad had superior throughput with 16 or 32 CPUs (Fig. S6A-B). In all cases, however, geNomad was most memory-efficient, likely due to Cenote-Taker 3 using pyhmmer for heavy computation whereas geNomad does not (Fig. S6C).

Cenote-Taker 3’s parallel scaling efficiency falls off above 4 CPUs and falls more steeply than geNomad (Fig. S6D). Therefore, we can recommend that users wishing to run Cenote-Taker 3 on many datasets deploy compute nodes with 4 CPUs for optimal efficiency.

**Figure S6.**
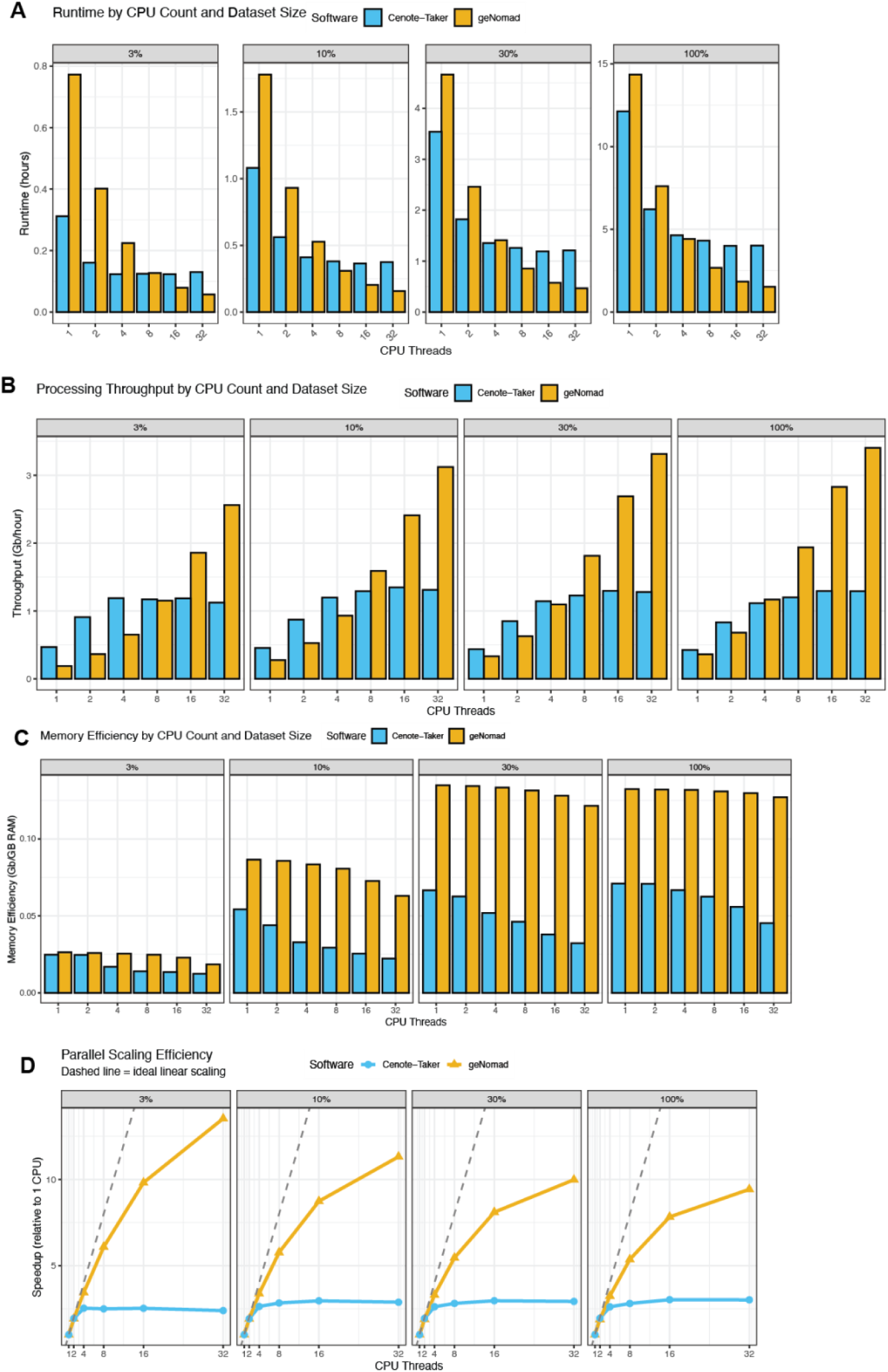
Performance Scaling for Virus Discovery. Three large metagenomic read sets (SRA: DRR582205, ERR9769281,ERR10905741) were assembled and these assemblies were combined to make an extremely large “stress test” for Cenote-Taker 3 and geNomad. The 100% dataset is 5.18 Gb of contigs; the 30% subset is 1.50 Gb; the 10% subset is 0.49 Gb; the 3% subset is 0.15 Gb. These different subsets were fed to Cenote-Taker 3 and geNomad with different resource allowances. (A) Runtime by CPU count and dataset size. (B) Throughput (Gb/hour) with different resource allowances. (C) Memory efficiency (Gb per Gigabyte of max RAM) with different resource allowances. (D) Parallel scaling efficiency.

### Other Features: Taxonomy, Prophage Extraction

Cenote-Taker 3 also assigns hierarchical taxonomy of virus contigs based on identity of its “hallmark” genes to GenBank virus records. A comparison of the aforementioned UHGV dataset of 100 high-quality MAGs shows very similar output to geNomad (Fig. S7A). Next, thousands of random GenBank genomes with NCBI taxonomy labels were downloaded, and Cenote-Taker 3 was used to annotate these to determine taxonomical concordance. For genomes where Cenote-Taker 3 detects a marker gene, we see >89% agreement with GenBank down to the family level (Fig. S7B). Cenote-Taker 3 will only classify genomes for which a hallmark gene is detected. When considering all contigs from this set, the accuracy drops to 73% at the family level (Fig. S7C), mostly due to the many reverse-transcribing viruses in GenBank, which Cenote-Taker 3 DB does not contain hallmark genes for, and small segments of segmented viruses.

Prophage extraction from bacterial chromosomes is performed by finding regions with virus “hallmark” genes and scoring surrounding genes for virus vs bacterial features and clipping the prophage where high bacterial gene content begins. Cenote-Taker 3 was compared to geNomad, VirSorter 2, and VIBRANT in a study by Wirbel, Hickey et al.^22^ for prophage boundary prediction accuracy. This study used soft clipping information to find prophage excision/integration points in bacterial MAGs and compared these coordinates to the boundaries predicted by the software. Overall, geNomad had the lowest error, and Cenote-Taker 3 had middle of the road performance. Cenote-Taker 3 boundary error was on par with geNomad after CheckV post-processing, suggesting a practical approach for Cenote-Taker 3-based pipelines.

**Figure S7.**
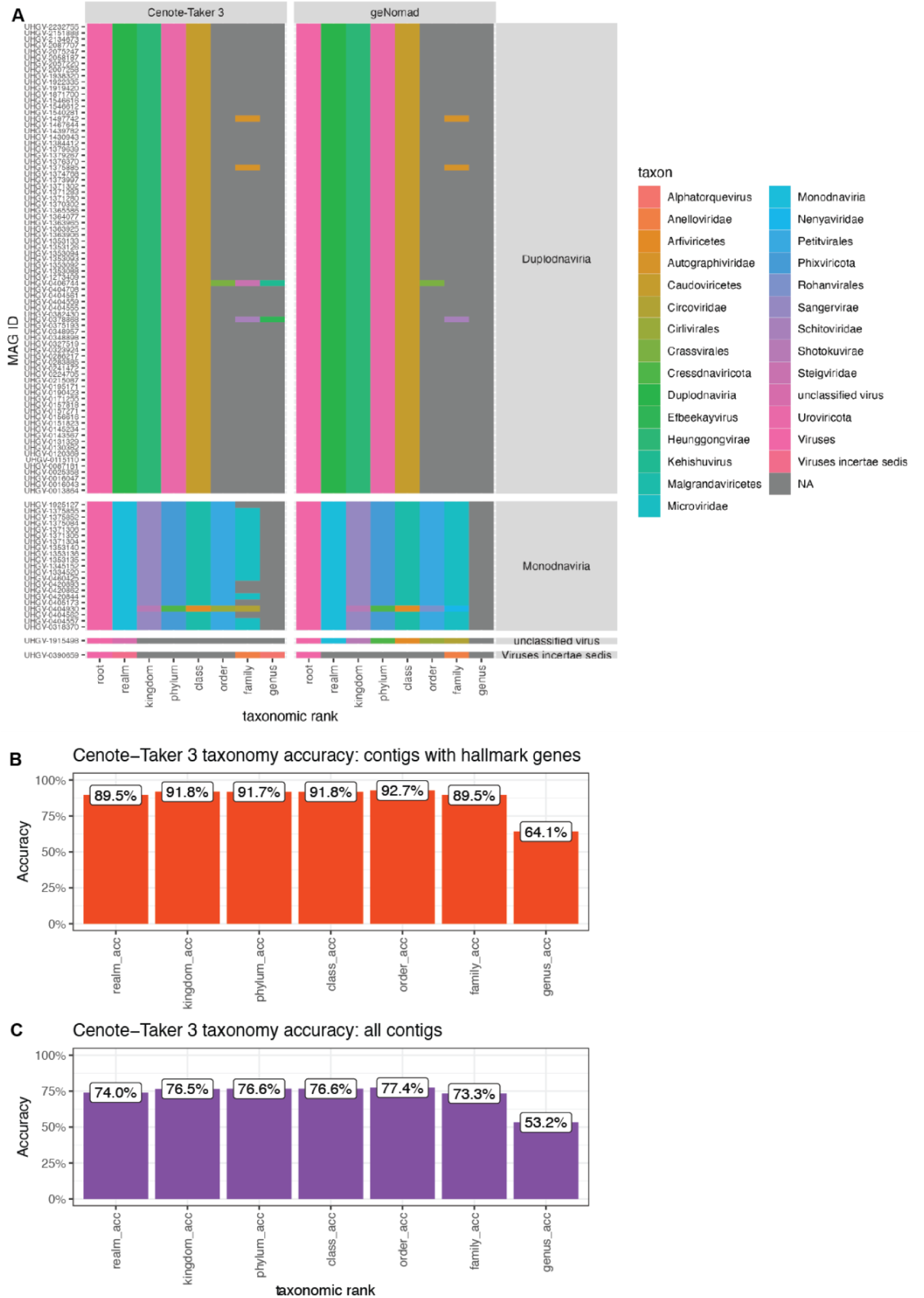
Taxonomic Benchmarking. (A) 100 high-quality virus MAGs from the UHGV dataset were run through Cenote-Taker 3 and geNomad and the hierarchical taxonomy labels were compared. Genomes were faceted by realm assigned by Cenote-Taker 3. (B) Random NCBI Virus genomes (n=2,959 contigs) in which Cenote-Taker 3 discovered hallmark genes were taxonomically classified with Cenote-Taker 3 and these hierarchical were compared to labels in official NCBI records. (C) Like B but with full set of random NCBI virus genomes (n=3,696). Cenote-Taker 3 will not assign taxonomy without discovering hallmark genes.

## Discussion

Cenote-Taker 3 is an ideal piece of software for annotation and cataloging of the virome, particularly if the focus is on high-quality genomes. It readily scales from single virus genomes to thousands of metagenomes. It can be used as an end-to-end program to identify virus MAGs, annotate their genes, excise prophage sequences from bacterial genomes, and taxonomically label virus sequences. Alternatively, it can also be used to simply annotate genes and report taxonomy of pre-discovered virus MAGs.

Cenote-Taker 3 reliably reports on the function of important genes and identifies viruses in complex datasets. Some existing software outperforms Cenote-Taker 3 in some ways, but this tool stands out for its combination of performance and accuracy. For example, the benchmarking performed showed that Cenote-Taker 3 annotated essential genes (MCP, TerL, portal protein for head-tail phages; RdRP and Capsid/Coat for RNA viruses) more accurately than other tools and finished annotation tasks more quickly than most tools (Fig. 2, Fig. S2-3). Virus discovery tests showed that Cenote-Taker 3 identifies some highly convincing virus MAGs that are missed by the field standard geNomad (Fig. 4-5). Therefore, we believe it’s justified to say that Cenote-Taker 3 occupies an impactful niche in viromics workflows, excelling at 1) functional annotation of previously uncharacterized virus genomes and 2) virome catalog building with a focus on complete/near complete MAGs. Cenote-Taker 3 will be essential for virome cataloging projects alongside other software, such as geNomad, used in this manuscript’s benchmarking.

Comparative tests for annotation and discovery of viruses, such as those performed in this study, also present discrete and useful opportunities for all software packages compared to improve. For example, when other software identifies a major capsid protein gene that Cenote-Taker 3 does not, this signal can be validated (e.g. using structural prediction) and the Cenote-Taker 3 database can be improved accordingly.

As long-read sequencing technologies continue to mature and become more affordable, tools like Cenote-Taker 3 will become increasingly valuable for producing high-quality complete viral genome annotations from metagenomic data. Future developments will focus on expanding reference databases to improve annotation of currently unknown genes, particularly those from under-sampled environments and viral lineages.

By offering simplified installation through Bioconda along with high performance on standard consumer hardware and high-performance compute systems, Cenote-Taker 3 contributes to democratizing viral genomics research, enabling labs with limited computational resources to conduct sophisticated analyses.

## Methods

### Cenote-Taker 3 Code Details

Cenote-Taker 3 is a command-line interface tool coded using Python and Bash. It imports and/or calls several bioinformatics packages. To read and write sequencing records, Biopython^23^ and Seqkit^24^ are used. To predict open-reading frames, pyrodigal-gv^5^ is used by default, and pyrodigal and phanotate^6^ are also available per command-line argument. Gene functional annotation is performed using pyhmmer^6^ and mmseqs2^18^. Hhsuite^25^ and its databases can also be used. trnascan-se^26^ is used to predict tRNA elements. Bedtools^27^ is used to deal with genomic ranges and coordinates. Minimap2^28^ and samtools^29^ are used for optional read alignment and quantification steps.

All code is publicly available and maintained on GitHub (https://github.com/mtisza1/Cenote-Taker3). The repo is also automatically backed up in Zenodo located at DOI:10.5281/zenodo.17290322.

Basic installation instructions using the mamba/conda package manager:

**mamba create -n ct3_env bioconda::cenote-taker3**

Bioconda package page can be accessed [here](https://bioconda.github.io/recipes/cenote-taker3/README.html#package-cenote-taker3). Quay container image, generated automatically by Bioconda, can be accessed [here](https://quay.io/repository/biocontainers/cenote-taker3).

### Cenote-Taker 3 Database Details

for the HMM databases, used for virus “hallmark” (virion, rdrp, and dnarep) gene detection as well as for some additional gene annotation tasks, the Cenote-Taker 2 database was expanded with gene models from PHROGs^30^, efams^31^, sequences from the Buck et al. survey^32^, and manually curated models. All model names were manually checked and updated as necessary to ensure similar models returned the same annotation. Notably, Cenote-Taker 3 hallmark databases do not contain gene models for reverse transcribing viruses (class Revtraviricetes). This is a design choice made to avoid overloading of results with endogenous retroviruses of eukaryotic genomes.

For the taxonomy database, NCBI’s NR Clustered cd90 amino acid sequence [database](https://ftp.ncbi.nlm.nih.gov/blast/db/experimental/) was downloaded on December 12, 2023, and filtered for records belonging the “Viruses” taxon. Then, these records were queried against the Cenote-Taker 3 hallmark databases (virion, rdrp, and dnarep) (DB v3.1.1), and hits were returned by standard Cenote-Taker 3 cut-offs. Hierarchical taxonomy labels were retrieved from GenBank for these selected records using taxonkit^33^. Only records which contained class, family, and genus labels were kept for the final taxonomy database. Mmseqs2 was used to create a taxDB with these records.

Database files can be accessed on [Zenodo](https://zenodo.org/records/12707420) (DOI 10.5281/zenodo.8429308)

### Annotation Benchmarks

Annotation benchmarks were performed using a 2023 MacBook Pro laptop computer with 16 GB of memory and an Apple M2 Pro chip running MacOS. Resource scaling benchmarks were performed on compute nodes with Intel “Emerald Rapids” with 32 central processing units and large onboard cache with 1 TB memory each.

The UHGV, Gut Circular, and Seawater Circular, Seawater virus-like particle RNA virus MAG, GenBank, RefSeq, short-read assemblies, and large metagenome sequences are available at [Zenodo](https://zenodo.org/records/16807783). Seawater virus-like particle RNA sequencing reads and assemblies are available on NCBI under bioproject PRJNA605028.

The following software package versions and databases were used: geNomad v1.11.1 (DB v1.9), MetaCerberus v1.4.0 (DB v1.4), Pharokka v1.7.3 (DB v1.4.0), phold v0.2.0 (DB v2), Cenote-Taker 3 v3.4.1 (DB v3.11).

All annotation software tested used pyrodigal-gv to conduct open reading frame prediction, making annotation rates as comparable as possible. The most appropriate setting for annotation were used based on software documentation. These are the threshold used by each tool, pharokka hmmer e-value <= 1e^-5^, MetaCerberus hmmer e-value <=1e^-9^, Cenote-Taker 3 hmmer e-value 1e^-7^ and mmseqs search e-value <= 1e^-3^, genomad mmseqs search <= 1e^-3^, phold foldseek e-value 1e^-3^. Example commands are as follows:

**genomad annotate --splits 4 seqs.fna gnmd_test1_out genomad_dbs/**
**metacerberus.py --prodigalgv seqs.fna --hmm “ALL” --dir_out metc_test1_out**
**pharokka.py -m -s -g prodigal-gv --meta_hmm -d pharokka_db -i seqs.fna -o phar_test1_out**
**phold run -i seqs.fna -o phold_test1_out**
**cenotetaker3 -c seqs.fna -r ct3_test1_out -p F -am T**

All scripts to reproduce analyses of annotation outputs are available on GitHub (https://github.com/mtisza1/ct3_benchmarks), and data are available on Zenodo.

### Discovery Benchmarks

Circular MAG assemblies from hot spring reads (DRR290133) and anaerobic digester sample reads (ERR10905741) are available on [Zenodo](https://zenodo.org/records/16807783). Reads were downloaded from SRA and assembled with myloasm v0.1.0 with default settings. Circularity of contigs is reported in myloasm output sequence headers.

GeNomad end-to-end pipeline was run on these data with default settings plus the “--enable-score-calibration” flag. Cenote-Taker 3 was run with the flags “-p T --lin_minimum_hallmark_genes 2 --circ_minimum_hallmark_genes 2” as recommended on the GitHub README for whole genome shotgun metagenome data.

Tool-specific contigs were annotated with phold v0.2.0 using default settings and visualized with Rstats package gggenomes^34^.

## Author Contributions Statement

Conceptualization: M.J.T., S.J.C., J.F.P.

Methodology: M.J.T., A.V.

Investigation: M.J.T., A.V.

Visualization: M.J.T.

Funding acquisition: S.J.C., J.F.P.

Project administration: S.J.C.

Supervision: J.F.P., S.J.C.

Writing—original draft: M.J.T., S.J.C.

Writing—review and editing: M.J.T., S.J.C., J.F.P., A.V.

## Acknowledgements

We are grateful all insights and suggestions from early users of this software, especially Dr. Chris Buck.

